# Implicit learning and exploitation of regularities involve hippocampal and prefrontal theta activity

**DOI:** 10.1101/611129

**Authors:** Eelke Spaak, Floris P. de Lange

**Affiliations:** Radboud University Nijmegen, Donders Institute for Brain, Cognition and Behaviour, Kapittelweg 29, 6525 EN, Nijmegen, The Netherlands

**Author notes:** Corresponding author: Eelke Spaak.

## Abstract

Observers rapidly and seemingly automatically learn to predict where to expect relevant items when those items are repeatedly presented in the same spatial context. This form of statistical learning in visual search has been studied extensively using a paradigm known as contextual cueing. The neural mechanisms underlying the learning and exploiting of such regularities remain unclear. We sought to elucidate these by examining behaviour and recording neural activity using magneto-encephalography (MEG) while observers were implicitly acquiring and exploiting statistical regularities. Computational modelling of behavioural data suggested that after repeated exposures to a spatial context, participants’ behaviour was marked by an abrupt switch to an exploitation strategy of the learnt regularities. MEG recordings showed that the initial learning phase was associated with larger hippocampal theta band activity for repeated scenes, while the subsequent exploitation phase showed larger prefrontal theta band activity for these repeated scenes. Strikingly, the behavioural benefit of repeated exposures to certain scenes was inversely related to explicit awareness of such repeats, demonstrating the implicit nature of the expectations acquired. This elucidates how theta activity in the hippocampus and prefrontal cortex underpins the implicit learning and exploitation of spatial statistical regularities to optimize visual search behaviour.

## Introduction

Objects are rarely encountered in isolation, but typically within certain spatial contexts. The encountered context, or scene, in which objects appear often induces clear expectations about where particular objects are likely to be located; e.g. we expect (and would more rapidly locate) a computer mouse next to the keyboard and not on top of the monitor. Such scene-based contextual expectations can greatly facilitate perception of individual elements of the scene (Biederman, 1972; Bar, 2004; Oliva and Torralba, 2007; De Lange et al., 2018). Humans are able to automatically and rapidly learn these spatial expectations through repeated exposure to the same or similar scenes, as demonstrated most forcefully by the experimental paradigm known as *contextual cueing* (Chun and Jiang, 1998; Chun, 2000; Goujon et al., 2015).

Contextual cueing is the effect robustly reported that observers engaged in visual search tasks become considerably faster in identifying target items in search displays that they have encountered before. Importantly, this effect becomes evident after only a few exposures to the same scene, and it appears to be fully implicit and automatic, happening outside of conscious awareness (Chun and Jiang, 1998). The robust nature and rapid establishment of the contextual cueing effect makes it ideally suited to study the neural and computational mechanisms involved in the acquisition of scene-based spatial expectations (Goujon et al., 2015).

Some work has been done on elucidating the neural mechanisms underlying this rapid learning and exploitation of statistical regularities, yet much remains unknown. Evidence from neuropsychology indicates that damage to the hippocampus and/or associated medial temporal lobe structures results in deficits in contextual cueing (Chun and Phelps, 1999; Manns and Squire, 2001), a finding which was later corroborated by functional magnetic resonance imaging (fMRI) (Greene et al., 2007; Giesbrecht et al., 2013). Additionally, fMRI studies have demonstrated the involvement in contextual cueing of parietal and frontal cortical structures associated with the top-down guidance of attention (Pollmann and Manginelli, 2009; Giesbrecht et al., 2013), lending support to the hypothesis that contextual cueing likely involves top-down guided attention based on a learned associative link. If, how, and when the hippocampal and parieto-frontal systems interact during learning and exploitation of regularities remains, however, unknown.

So far, little work has been done in understanding the role of neural oscillations in the acquisition and exploitation of the associative link underlying contextual cueing. This is surprising, considering the proposed role of especially low-frequency oscillations (theta/alpha/beta) in communicating predictive and top-down information within and across brain regions (Lisman and Jensen, 2013; Jensen et al., 2014; Kerkoerle et al., 2014; Bastos et al., 2015; Michalareas et al., 2016; Spaak et al., 2016; Chao et al., 2018). Given this proposed (and reported) role of neural oscillations, we used the contextual cueing paradigm to examine the neural oscillatory mechanisms involved in the acquisition and exploitation of spatial predictions.

The learning of an associative link (especially involving the hippocampus, which is crucial for declarative memory) and the top-down guidance of attention are often thought to be associated with deliberate, conscious thought (Eichenbaum, 2000; Dehaene et al., 2006; Cohen et al., 2012). Interestingly, the spatial predictions learnt in contextual cueing, possibly depending on the same mechanisms, are typically thought to be fully implicit and automatic. Many studies have made such a claim, based on an explicit recognition task following the main visual search task (e.g. (Chun and Jiang, 1998)) or more elaborate target-generation tasks (Chun and Jiang, 2003). However, it has been demonstrated that the statistical power to detect conscious awareness in these paradigms was limited, and a more powerful meta-analysis has shown that participants were likely aware of the contextual repeats to some extent (Vadillo et al., 2016). We therefore additionally set out to quantify to what extent the benefit derived from previous exposures to a scene depends on conscious awareness of these exposures or not.

To answer these questions, we recorded magnetoencephalography (MEG) data while participants were performing a variant of the classical contextual cueing search task. To preview our results: during the first few blocks, as participants were becoming familiar with the repeated scenes, we observed stronger hippocampal theta activity for familiar (old) compared to unfamiliar (new) scenes. During these blocks, however, there was no behavioural advantage yet for repeated scenes. After the first few blocks, there was a marked shift in both neural processing and behavioural performance. Now frontal cortex (but not hippocampus) showed elevated theta activity for old compared to new scenes, and participants showed a marked behavioural benefit for these familiar scenes. This benefit clearly occurred outside of conscious awareness of scene familiarity; in fact, larger awareness was associated with a smaller benefit. These findings thus shed light on how the brain rapidly extracts statistical regularities from our environment and can use them to guide goal-directed behaviour, even without conscious awareness.

## Results

Human volunteers (N = 36) participated in a variant of the classical contextual cueing task. Participants were instructed to locate a target T stimulus amongst L-shaped distractors, and report the orientation of the T (tilted clockwise/counterclockwise) with a button press (Figure 1A). The experiment consisted of 22 blocks, and in each block the same 20 familiar displays were presented (‘Old’ trials) in random order, randomly intermixed with 20 unfamiliar displays (‘New’ trials). In later blocks, we additionally included a small amount of violation trials to be able to study the effects of different types of violations of the established contextual predictions. From block 9 onwards, 5 trials per block were changed from an Old trial into a ‘Target violation’ trial by exchanging the target location with a randomly chosen distractor (thus disrupting the context–target association), and 5 trials per block were changed from an Old trial into a ‘Distractor violation’ trial by randomly rotating each distractor a random multiple of 90° (yet preserving the global context–target association) (Figure 1B).

**Figure 1:**
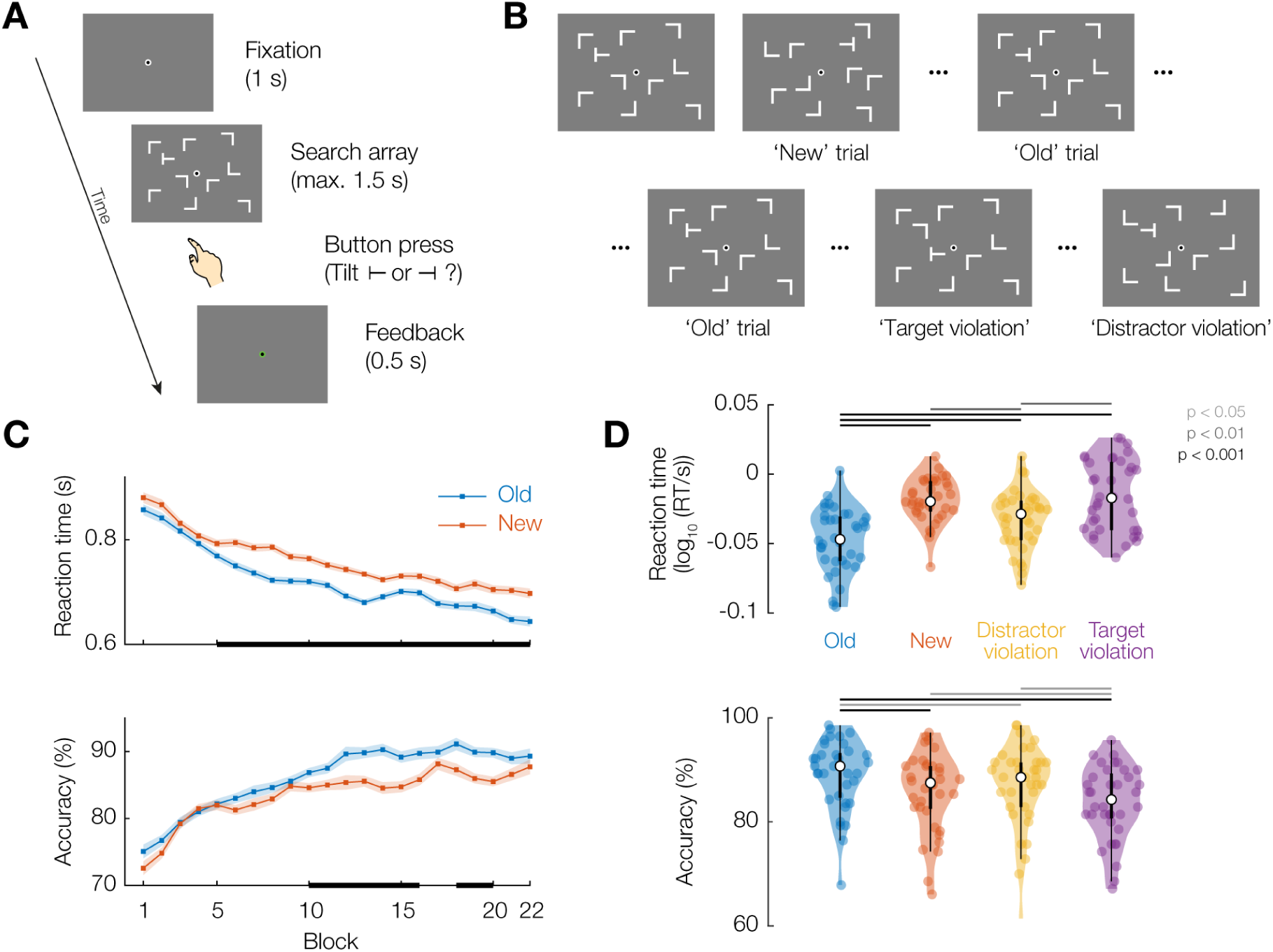
Experimental design and search task behavioural results. **(A)** Schematic of an individual search task trial. **(B)** Different trial types: ‘New’ trials are unfamiliar; ‘Old’ trials are repeated every block; on ‘Target violation’ trials the target on an Old display swaps place with a random distractor; ‘Distractor violation’ trials maintain the spatial configuration of an Old display, but every distractor is rotated a random multiple of 90°. **(C)** Reaction time and accuracy time courses over the experiment for Old and New trials (error shading indicates within-participant corrected standard error of the mean). Black bars on the x-axis indicate pointwise t-tests p < 0.05 for illustration. **(D)** Reaction time and accuracy for the four conditions, averaged over blocks 9–22 (well after learning has occurred). Coloured dots are individual participants. Inside the violin plot, the white dot reflects the median, thick box indicates quartiles, thin line indicates quartiles ± 1.5 × inter-quartile range.

### Search task behaviour demonstrates robust contextual cueing

Over the course of the experiment, responses became faster and accuracy increased (Figure 1C). During the later part of the experiment (*a priori* defined as blocks 9–22), participants showed a clear reaction time benefit for Old displays (678 ± 91 ms; mean ± s.d. across participants), compared to New ones (727 ± 82 ms; t_35_ = 6.59, p = 1.2 × 10^−7^, BF_10_ = 1.1 × 10^5^), as well as a higher accuracy (Old: 89 ± 7 %; New: 86 ± 8 %; t_35_ = 5.30, p = 6.4 × 10^−6^, BF_10_ = 3.1 × 10^3^) (Figure 1D). We thus replicate the classical contextual cueing benefit for visual search in repeated displays.

Responses on Distractor violation trials were slower (710 ± 100 ms) than on Old trials (t_35_ = 5.61, p = 2.5 × 10^−6^, BF_10_ = 7.4 × 10^3^), and slightly less accurate (87 ± 8 %; t_35_ = 2.32, p = 0.03, BF_10_ = 1.9); but faster than on New trials (t_35_ = 3.01, p = 0.0048, BF_10_ = 7.9), with no difference in accuracy (t_35_ = 1.18, p = 0.3, BF_10_ = 0.34) (Figure 1D). This indicates that the contextual cueing effect is, at least partly, due to the global configuration of the display, and not exclusively to its local features.

Reaction times on Target violation trials (735 ± 105 ms) were slower than on Old trials (t_35_ = 5.86, p = 1.2 × 10^−6^, BF_10_ = 1.5 × 10^5^), slower than on Distractor violation trials (t= 3.05, p = 0.004, BF_10_ = 8.6), and not distinguishable from New trials (t_35_ = 0.678, p =0.50, BF_10_ = 0.22). Accuracy on Target violation trials (84 ± 7 %) was substantially lower than on Old trials (t_35_ = 6.22, p = 3.9 × 10^−7^, BF_10_ = 4.1 × 10^5^), and also somewhat lower than on New (t_35_ = 2.55, p = 0.015, BF_10_ = 3.0) and Distractor violation (t_35_ = 2.54, p = 0.016, BF_10_ = 2.9) trials (Figure 1D). This indicates a contextual cost when the spatial prediction learned from repeated exposures is violated.

### Explicit recognition is inversely related to implicit behavioural benefit

After completion of the main search task, participants were asked whether they had the “feeling that some of the search displays occurred multiple times over the course of the experiment” and responded with a button press to indicate ‘Yes’ or’ ‘No’. Subsequently, participants completed a final (unspeeded) block of 20 New and 20 Old search displays, where the task was to indicate whether they had seen that particular display before during the main task. This allowed us to assess participants’ subjective feeling of recognition (question), their objective explicit memory for the search arrays (recognition task), and their relationship to each other and the contextual cueing effect. Results for these analyses are depicted in Figure 2.

**Figure 2:**
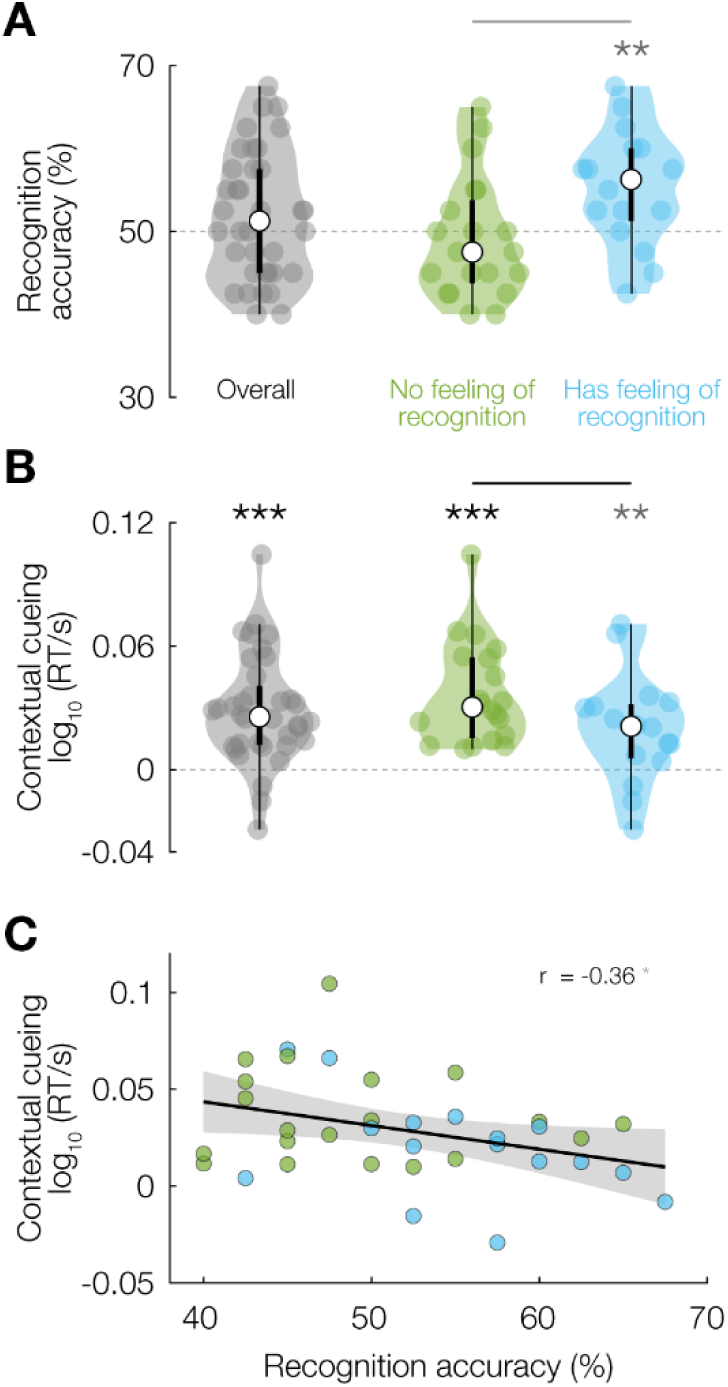
Recognition task behavioural results. **(A)** Accuracy in recognizing search displays as Old or New during the recognition task; for the whole sample (left, grey), and split by whether participants indicated a feeling of repeated displays during the main task (middle, green) or not (right, blue). **(B)** Contextual cueing effect during the main search task (defined as RT difference between Old and New trials), for the same groups as in (A). **(C)** Contextual cueing effect during the main task as a function of recognition performance. Dots are individual participants, coloured by their response to the feeling-of-recognition question. * p < 0.05, ** p < 0.01, *** p < 0.001. Significance bars and violin plot details as in Figure 1.

As commonly reported in studies of contextual cueing in visual search, the experimental sample as a whole was at chance (50 %) in distinguishing Old from New displays during the recognition task (52 ± 8 %; t-test versus chance level t_35_ = 1.46, p = 0.15), though the data were only ∼2 times more likely under the null hypothesis of chance level than under the alternative hypothesis (BF_10_ = 0.47). 16/36 participants (44 %) reported having the feeling that some displays were repeated, while 20/36 (56 %) reported having no such feeling. Interestingly, participants who felt that some of the displays were repeated were on average also slightly above chance level in recognizing displays as Old during the recognition task (55 ± 7 %; t_15_ = 3.03, p = 0.0084, BF_10_ = 6.33). In contrast, participants who did not have the impression that displays had been repeated were at chance level in recognizing the displays (49 ± 7 %; t_19_ = −0.538, p = 0.60, BF_10_ = 0.27) (Figure 2A). There was thus substantial variability in objective explicit recognition performance among participants, which furthermore appears directly related to the subjective feeling of having encountered displays before.

Of considerable interest is the relationship we observed between the subjective feeling of having observed repeats and the strength of the contextual cueing effect during the main search task (Figure 2B). There was a clear reaction time benefit for Old versus New displays in both the Recognizing group (i.e., those who answered ‘Yes’ to the feeling-of-repeats question; t_15_ = 3.02, p = 0.0087, BF_10_ = 6.1) and the Non-recognizing group (i.e., those who answered ‘No’; t_19_ = 6.58, p = 2.7 × 10^−6^, BF_10_ = 7.3 × 10^3^). Surprisingly, the contextual cueing effect was considerably larger in the Non-recognizing group (t_34_ = 24.6, p < 10^−16^, BF_10_ = 5.9 × 10^19^) (Figure 2B), suggesting that search benefits were larger when participants were not aware of the regularities. Further supporting this notion, individual participants’ accuracy on the recognition task was negatively correlated with the contextual cueing effect during the main task (r = −0.36, t_34_ = −2.09, p = 0.044), although the evidence for this relationship was anecdotal (BF_10_ = 1.3) (Figure 2C). This relationship between both the objective explicit recognition performance and the subjective feeling-of-repeats on the one hand, and the contextual cueing benefit during the main search task on the other, is rather striking: individuals with *less* explicit knowledge of context-target associations show a *greater* behavioural benefit when exploiting such associations.

### The contextual cueing benefit appears suddenly and not gradually

The curves of reaction time for Old and New trials over the course of the experiment (Figure 1C) appear to diverge from about block 5 onwards, after which the difference between them appears to remain stable. We sought to quantify the evidence for this potentially interesting descriptive observation through Bayesian model comparison. Specifically, per participant, we fitted five models to the condition-specific reaction time data: No effect (no difference between conditions), No switchpoint (constant difference between conditions), Switchpoint (no difference between conditions until a particular point in time in the experiment and a constant difference afterwards), Linear (linearly increasing difference, trialwise across the experiment time), and Blockwise linear (linearly increasing difference, blockwise across the experiment time) (Figure 3A).

**Figure 3:**
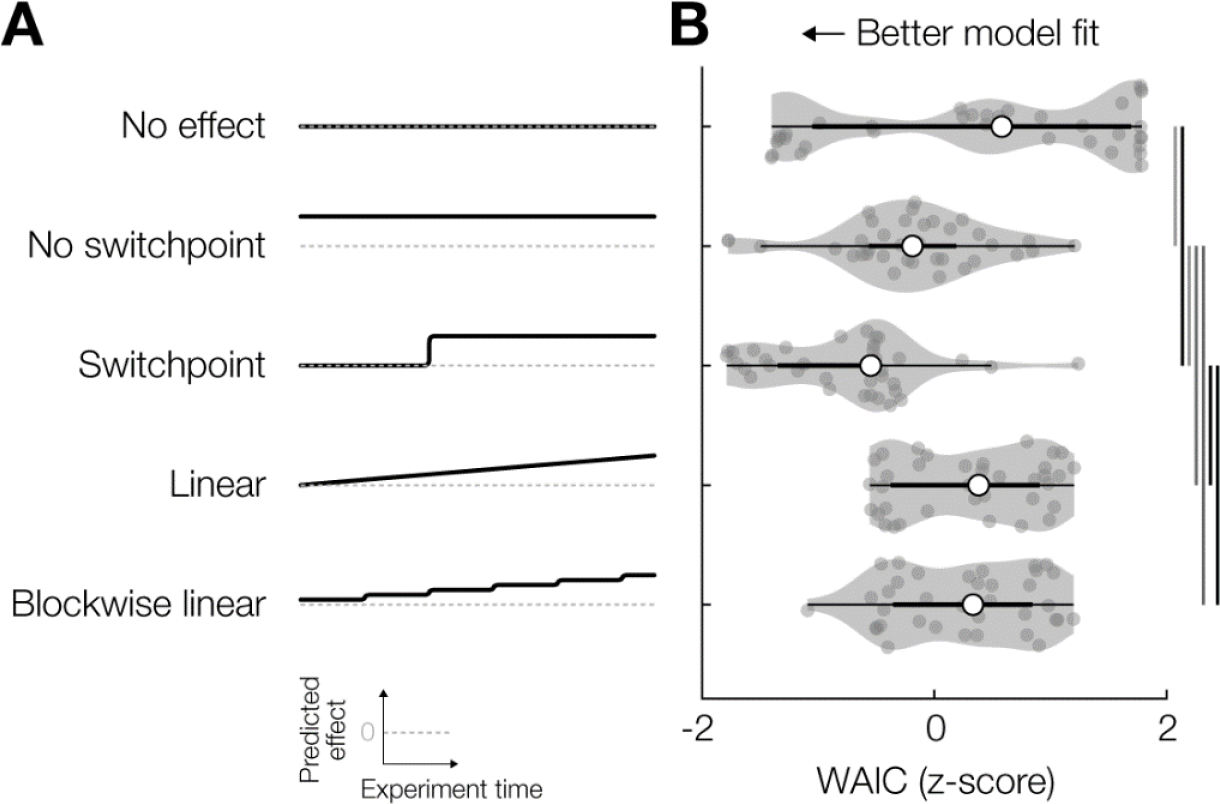
Reaction time modelling results. **(A)** Illustration of the five different models that were fit to the reaction time data for Old and New trials. Grey dashed line indicates zero in each subpanel; black solid line indicates the predicted evolution of the contextual cueing effect (RT_New_-RT_Old_) over the course of the experiment. **(B)** Watanabe-Akaike Information Criterion scores (z-scored across models per participant) for each of the models. Significance bars and violin plot details as in Figure 1.

Across our experimental sample, we observed the strongest evidence (quantified by Watanabe-Akaike Information Criterion, WAIC) for the Switchpoint model (relative WAIC = −288.04), followed by the No switchpoint (WAIC = −227.87), Blockwise linear (WAIC = −207.55), Linear (WAIC = −206.34), and No effect (WAIC = −36.99) models. The pairwise differences between Switchpoint and all the other models were significant and substantial (versus No switchpoint: t_35_ = 2.71, p = 0.010, BF_10_ = 4.1; versus Blockwise linear: t_35_ = 6.08, p = 6.1 × 10^−7^, BF_10_ = 2.8 × 10^4^; versus Linear: t_35_ = 3.44, p = 0.0015, BF_10_ = 21; versus No effect: t_35_ = 5.03, p = 1.5 × 10^−5^, BF_10_ = 1.4 × 10^3^) (Figure 3B). This indicates that, indeed, observers first acquire the contextual knowledge (implicitly), and then switch to a strategy in which they start benefitting from repeated exposures to the same display.

### Learning a predictive context is associated with hippocampal theta activity

During the experiment, we continuously recorded magnetoencephalography (MEG) data from our participants, allowing us to characterize the neural dynamics during the learning and exploitation of the associative link between scene context and target location. We were specifically interested in the role of low-frequency neuronal oscillations in contextual cueing, with repeated versus novel spatial contexts (i.e., Old versus New trials) as the main contrast of interest. We computed time-frequency representations of oscillatory power, averaged over all trials per condition in the experiment, and contrasted these values between conditions. The data were significantly different between Old and New (cluster-based permutation test p = 0.004), with a prominent difference during the first 500 ms of stimulus processing in the theta^1^ frequency band (1–7 Hz). The MEG sensor topography of this difference was rather diffuse, with local left frontal/central and right anterior temporal peaks (Figure 4A).

**Figure 4:**
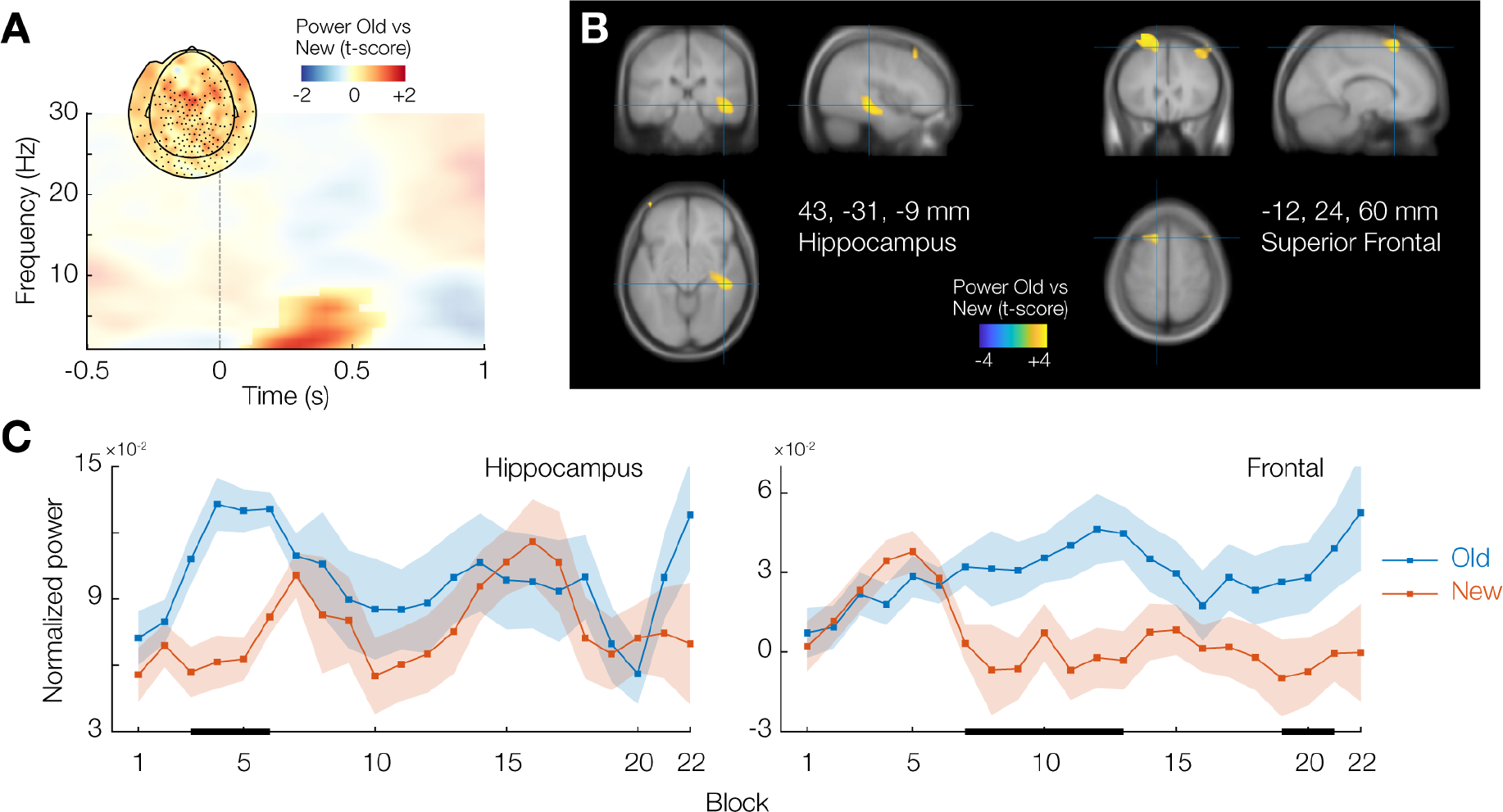
Theta band activity differences between Old and New trials. **(A)** Time-frequency representation of sensor-level power difference between Old and New trials across the whole experiment. Opacity mask indicates cluster-corrected significant differences (p < 0.05) for illustration. Inset shows the sensor topography of the effect; significantly contributing sensors are highlighted with a dot. **(B)** Source-level contrast of the time-frequency window of interest identified in (A), projected on the average Montreal Neurological Institute (MNI) template brain. MNI coordinates given are of two local maxima; anatomical labels are from the Automated Anatomical Labeling (AAL) atlas. **(C)** Source-level power values for Old and New trials separately, for the two identified regions in (B), as a function of experiment time.

To get a clearer view of the neural origin of these effects, we performed beamformer source analysis of the theta activity (1–7 Hz) in the time window 0–500 ms post stimulus onset, and again computed condition contrasts. The strongest source of this effect was located in the right hippocampus (peak at 43, −31, −9 mm in Montreal Neurological Institute (MNI) space; Figure 4B, left panel).

We next investigated the evolution over the course of the experiment of the difference in stimulus-related hippocampal theta activity between Old and New trials. Interestingly, this difference was primarily evident during blocks 3–6, but not during the later part of the experiment (Figure 4C, left panel). For the majority of this time period, a behavioural contextual cueing effect was not yet evident (Figures 1C, 5A). This suggests that the hippocampal theta rhythm is specifically associated with the *learning* of the predictive link between spatial context and target location, and that, once learned, hippocampal theta activity returns to baseline levels.

**Figure 5:**
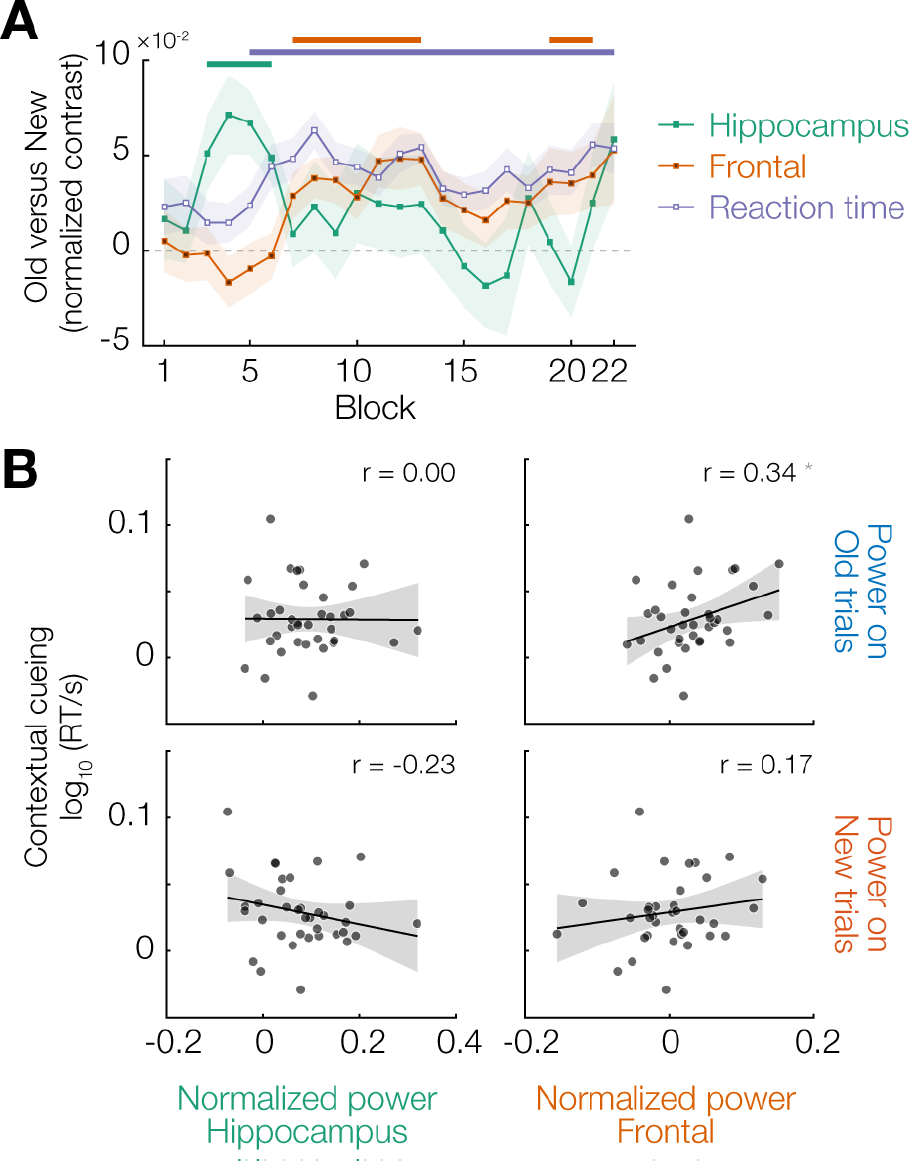
Relationship between neural and behavioural modulation. **(A)** Evolution of theta power differences (Old - New) over the course of the experiment in the right hippocampus and frontal cortex clusters identified in Figure 4, as well as the contextual cueing reaction time effect (New - Old). **(B)** Scatter plots of the behavioural contextual cueing effect versus theta power in the two identified clusters, separately for Old and New trials. Each dot is a participant; black lines are the best-fitting linear regression, with grey shading indicating the 95% confidence interval.

### Exploiting a predictive context is associated with frontal theta activity

In addition to the peak in right hippocampus, several sources in frontal cortex also contributed to the difference in theta power between Old and New trials. The strongest frontal contribution was from the left superior frontal cortex (MNI x, y, z = −12, 24, 64 mm; Figure 4B, right panel), with additional, weaker, local maxima in right middle frontal gyrus (44, 24, 48 mm) and left precentral gyrus (−44, 0, 40 mm).

Interestingly, the evolution of this frontal effect over the time course of the experiment was markedly different from that observed in the hippocampus. Theta power for Old and New trials started to diverge significantly from block 7 onwards, and remained distinct throughout the experiment (Figure 4C, right panel). The frontal theta effect thus emerged as the hippocampal effect was ramping down, and closely mirrored the behavioural benefit for reaction times (Figure 5A).

Although frontal theta was clearly different between Old and New trials, there were no significant pairwise differences between either of the violation conditions (Distractor violation or Target violation) and Old or New (all p > 0.05), with evidence for the null hypothesis of no difference ranging from moderate to inconclusive (0.19 < BF_10_ < 0.88). We therefore cannot exclude the possibility that the absence of an effect here is due to the substantially lower trial count for the violation conditions (65 per condition for each violation condition, versus 355 for Old).

The time courses in Figure 5A suggest that the reaction time benefit is primarily driven by theta activity in the frontal cortex, presumably reflecting top-down driven guidance of attention based on context-induced predictions about target location. If this is the case, then theta power should correlate with the reaction time benefit specifically on Old trials (and not on New, since no predictive link was learned for these), and specifically in the frontal cortex (and not the hippocampus). This is indeed what we found: frontal theta power on Old trials was significantly correlated across participants with the contextual cueing effect (r = 0.34, t_34_ = 2.13, p = 0.041), but there was no such relationship for the hippocampus or for New trials (all p > 0.10, 0.18 < BF_10_ < 0.45) (Figure 5B). However, we note that the evidence for the correlation between frontal theta and the contextual cueing effect, although significant, was inconclusive (BF_10_ = 1.3), and should thus be interpreted with caution.

### Neither hippocampal nor frontal theta is associated with explicit recognition

As mentioned, we observed substantial variability among participants in explicit recognition performance, and a striking inverse relationship with the behavioural contextual cueing benefit (Figure 2). An interesting possibility therefore would be that either the hippocampal or the frontal theta effect dissociates between recognizers and non-recognizers. However, we found neither to be the case. We found no evidence that frontal theta in blocks 9–22 was different between the Recognizing and Non-recognizing groups (t_34_ = 1.69, p = 0.10, BF_10_ = 0.97), and frontal theta was not correlated with recognition performance (r = 0.11, t_34_ = 0.65, p = 0.52, BF_10_ = 0.22). We also found no evidence that hippocampal theta in blocks 3–6 was different between the Recognizing and Non-recognizing groups (t_34_ = 1.47, p = 0.15, BF_10_ = 0.75), or that it was correlated with recognition performance (r = 0.29, t_34_ = 1.79, p = 0.08, BF_10_ = 0.76). The theta band effects described above are thus specifically related to the implicit learning and behavioural exploitation of scene-based spatial predictions, and are unlikely to be explained by either the recognizers or the non-recognizers showing the effects, and not the other.

## Discussion

We set out to elucidate the neural oscillatory mechanisms underlying the acquisition and exploitation of predictive links between global scene context and search target location. Repeated displays were associated with elevated hippocampal theta power early in the experiment, while there was no behavioural benefit yet. Strikingly, as the hippocampal activity difference receded, stronger frontal theta activity became apparent for repeated scenes, concomitantly with a clear reaction time benefit. This reaction time benefit was furthermore positively correlated with frontal theta power for repeated scenes, and negatively related to participants’ explicit recognition of repeated search displays. A hippocampal-prefrontal interplay in the theta band thus underlies the acquisition and exploitation of unconscious scene-based spatial predictions, demonstrating a potential mechanism through which perception can be facilitated by prior expectations (Biederman, 1972; Bar, 2004; Oliva and Torralba, 2007; Summerfield and de Lange, 2014; de Lange et al., 2018).

### Contextual cueing is driven by overall configuration, implicit in nature, and all-or-none

It has been claimed that scene-based expectations are generated based on a rapid extraction of low spatial frequency information from an encountered scene, which can subsequently guide more fine-grained explorations (Bar, 2004). The Distractor violation condition allowed us to investigate to what extent the contextual cueing effect depends on local or global aspects of the search displays. Rotating individual distractors preserves the low spatial frequency distribution of the display, but perturbs the location of individual high spatial frequency edges. If global features, and thus low spatial frequency content, are indeed exclusively responsible for contextual cueing, then Distractor violations should be processed as efficiently as Old displays. We found that Distractor violations are instead processed less efficiently than Old displays (slower RT, slightly lower accuracy), yet still considerably more efficient (faster and more accurate) than New displays. Overall configuration thus indeed seems critical for the contextual cueing effect, though not exclusively responsible for it. This is in line with previous findings (Chun and Jiang, 1998; Jiang and Wagner, 2004).

Our experimental sample of participants was at chance level in recognizing individual displays as Old or New after the main search task. This is a common finding in contextual cueing studies, and is interpreted as evidence that the familiarity after repeated search displays is a form of fully implicit knowledge (Chun and Jiang, 2003; Goujon et al., 2015). The implicit nature of this effect has been called into question, however, because individual studies have lacked the statistical power to detect above-chance performance (Vadillo et al., 2016). Also in our case, the evidence for the null hypothesis of chance level was only anecdotal across the whole sample, as indexed by Bayesian analysis. A more powerful meta-analysis has instead demonstrated evidence for a weakly *above*-chance recognition of search displays (Vadillo et al., 2016). In our experiment, we included a subjective ‘sense of repeated exposure’ question after the main search task, only after which the participants performed the recognition task. Some participants appeared to exhibit some awareness of the repeats (as indicated by the group answering that they felt that there were repeats indeed being above-chance at recognition performance), while some had no such awareness (substantial evidence for at-chance performance, no feeling of repeats). We found a considerably *larger* contextual cueing effect in the group *without* awareness, and a *negative* correlation of behavioural benefit with recognition performance. This demonstrates that the behavioural contextual cueing effect is, indeed, likely implicit in nature. Nonetheless, observers might additionally acquire explicit recognition knowledge of the search displays (thus explaining the above-chance result found in the meta-analysis by Vadillo et al.)

The inverse relationship between awareness of the regularities and the performance benefit derived from the regularities is in line with the idea that statistical learning biases sensory and decision processing in an automatic and non-deliberate fashion. Indeed, the benefits of statistical learning in visual search have previously been reported particularly when participants were instructed to “be as receptive as possible and let the [target] ‘pop’ into [their] mind”, but not when participants were instructed to “actively search for the target” (Mühlenen and Lleras, 2004). This corroborates the notion that contextual cueing appears grounded in automatic, unconscious, perception; or, in Kahneman’s terms, “System 1” thought (Kahneman, 2013); with any involvement of System 2 (deliberate, conscious thought) abolishing, or at least reducing, the effect.

A final result we observed in our behavioural data is that the contextual cueing effect likely emerges suddenly, and not gradually. To our knowledge, this has not been reported (or explicitly tested) before. This suggests that contextual cueing is a form of all-or-none learning (Jones, 1962; Estes, 1964), which has also been shown to underpin simple paired associate learning (Brainerd and Howe, 1978). The all-or-none nature of the phenomenon under study here ties nicely to the fact that synaptic long-term potentiation (LTP) is thought to be discrete and not continuous (Murthy, 1998). Synapses between hippocampal CA3 and CA1 subregions, specifically, are strengthened or weakened in an all-or-none manner (Petersen et al., 1998), which is especially interesting given the hippocampal effects we report in this study.

### Hippocampal and prefrontal theta activity underlies learning and exploitation of scene-based predictions

A wide body of evidence has linked the hippocampal theta rhythm to the encoding of memories, both spatial and non-spatial in nature (O’Keefe and Nadel, 1978; O’Keefe and Recce, 1993; Hasselmo, 2005; Buzsaki and Moser, 2013; Colgin, 2013, 2016; Lisman and Jensen, 2013; Staudigl and Hanslmayr, 2013; Backus et al., 2016; Bellmund et al., 2018). We now demonstrate, for the first time, that specifically the acquisition, and not the retrieval, of memories for spatial context in a visual search task involves the hippocampal theta rhythm. Although the involvement of the theta rhythm in contextual cueing has not been reported before, the hippocampus *per se* has been linked to contextual cueing through several strands of evidence. First, lesions to the hippocampus and/or associated medial temporal lobe structures result in severe deficits in the contextual cueing effect (Chun and Phelps, 1999; Manns and Squire, 2001). Second, BOLD activity in the right hippocampus is reduced for Old displays compared to New ones during the learning phase of a contextual cueing task (Giesbrecht et al., 2013). We observed an *increase* in hippocampal theta power for Old versus New displays, whereas Giesbrecht et al. reported a *decrease* in BOLD for Old versus New. This apparent discrepancy can be reconciled by noting that hippocampal theta and BOLD activity tend to be anti-correlated, specifically during memory encoding (Fellner et al., 2016). Our observation of theta activity specifically in the right, and not the left, hippocampus, is in line with the fact that the right, and not the left, hippocampus appears specifically involved in spatial memory (Burgess et al., 2002).

It has been debated whether MEG is sensitive enough to measure signals from deep brain structures, such as the hippocampus. In recent years, however, a consensus has emerged that, given proper source reconstruction techniques, MEG is indeed able to measure signals from the hippocampus (Dalal et al., 2013; Pu et al., 2018; Pizzo et al., 2019). We furthermore note that we did not start with an anatomical hippocampus-based region of interest and test the signals coming from that region, but instead used a data-driven approach which subsequently yielded the hippocampus as a strong and significant source of sensor-level effects. Considering these facts, together with the well-established role of the hippocampal theta rhythm in (spatial) memories (thus rendering the effect a priori plausible), we consider the hippocampus to be indeed a likely source of the observed sensor-level effects.

The exploitation of the learned link between spatial context and target location was associated with a clear theta frequency band effect in prefrontal cortex. The peak source of this effect was located in *superior* prefrontal cortex, which is known to be associated with the top-down guided orienting of (spatial) attention (Corbetta and Shulman, 2002). We believe attentional orienting to be the most likely explanation for the frontal theta effect reported here, since it has been shown that participants locate the target more efficiently in repeated displays due to rapid orienting of spatial attention (Zhao et al., 2012, 2013). Frontal theta activity is also commonly linked to the retrieval of memories (Kahana et al., 2001; Osipova et al., 2006; Addante et al., 2011), which might additionally partly explain the involvement of this rhythm in our experiment. Finally, it is interesting to note that several authors have suggested that these two phenomena (i.e., memory retrieval and attentional orienting) should be considered two sides of the same coin, (partly) dependent on the same neural mechanisms (Chun and Turk-Browne, 2007; Awh et al., 2012; Hutchinson and Turk-Browne, 2012).

Intriguingly, the two neural systems identified here (hippocampus during learning; frontal cortex during exploitation) to underlie clearly *implicit* memory are typically associated with *explicit* processes. The hippocampus is crucial for declarative memories, which are by definition amenable to conscious access (Eichenbaum, 2000). The top-down guided (i.e., endogenous) orienting of attention by frontal cortex is often thought to be intimately related to deliberate, conscious thought (Dehaene et al., 2006; Cohen et al., 2012) (although this relationship is fiercely debated, see Koch and Tsuchiya (2007)). We now demonstrate that learning and exploitation of fully implicit knowledge is associated with these two brain structures, and explicit awareness of the same knowledge is not. An interesting corollary is thus that activity in these regions is not necessarily associated with conscious awareness of what is being learned.

## Conclusion

Recent years have seen considerable interest in and support for the idea that the brain is essentially a prediction machine, continuously trying to minimize the mismatch between expectations and sensory inputs (Clark, 2013). The ubiquitous context effects in perception (Bar, 2004; Oliva and Torralba, 2007; de Lange et al., 2018), established through statistical learning (Turk-Browne et al., 2010), are thus explained by noting that perception depends strongly on prior expectations and not solely on sensory input. These prior expectations, in turn, are continuously fine-tuned in order to optimally process future input, and so on. We demonstrate that the acquisition and exploitation of scene-based expectations involve theta-band dynamics in an interplay between the hippocampus and the prefrontal cortex, which happens outside of conscious awareness. These results shed light on how humans are able to rapidly adapt their prior expectations in order to optimally guide perception and behaviour.

## Methods

### Participants

We recruited 36 healthy right-handed participants (23 female; 13 male; age 24 ± 5 years (mean ± standard deviation), range 17–45 years) from the Radboud University participant pool. The sample size was chosen to obtain ≥ 80 % power for detecting a medium effect size (d = 0.5) with a two-sided paired t-test at an alpha level of 0.05. All participants had normal or corrected-to-normal vision. Participants received € 12 compensation for their participation in the main MEG experiment, and an additional € 5 in case they returned for an anatomical magnetic resonance imaging (MRI) scan (i.e., if no such scan was available yet from participation in other experiments). The study was approved by the local ethics committee (CMO Arnhem-Nijmegen, Radboud University Medical Center) under the general ethical approval for the Donders Centre for Cognitive Neuroimaging (“Imaging Human Cognition”, CMO 2014/288). Participants provided written informed consent prior to the experiment.

### Stimuli, task, and experimental design

Throughout the experiment, a fixation dot (outer white diameter 8 pixels; inner black diameter 4 pixels) was presented in the center of the screen. Participants were asked to blink as little as possible and keep looking at the central fixation dot. Participants were instructed to search for a T-shaped target stimulus amongst L-shaped distractors (all stimuli measured 1.5 × 1.5 ° of visual angle and were coloured white, presented on a grey background). Distractor L shapes were rotated a random multiple of 90 °. The target T could be rotated 90 ° to the left or right, and participants made a left or right handed index finger button press to indicate the orientation of the T. Each trial started with a 1 s fixation period, and search displays were presented for 1.5 s or until the response button press. After the button press, participants were informed whether their response was correct or not by the white part of the fixation dot turning green (correct) or red (incorrect) for 0.5 s (Figure 1A). Search stimuli were arranged pseudorandomly on an 8 × 6 regular grid spanning −9 to +9 ° horizontal and −6 to +6 ° vertical from the center of the screen, with an added amount of random jitter ± 0.4 ° per stimulus to prevent exact collinearities (Chun and Jiang, 1998). The target was constrained to always appear between 6 ° and 8 ° of excentricity, and the mean distance between the target and all distractors was constrained to be between 9 ° and 11 °.

The main search task consisted of 22 blocks of 40 trials each. Participants were given the option to take a short break (while remaining seated in the MEG chair) every second block. We embedded repeats of search displays in each block to establish the classical contextual cueing effect, i.e. the ‘Old’ displays. The search configuration of an Old trial was exactly the same across blocks, i.e. the location and rotation of all distractors was maintained, as well as the location of the target. Importantly, the orientation of the target was always randomly chosen and not repeated, so participants could not learn associations between a given display and the appropriate response, only between the display and the target location. All search blocks contained 20 ‘New’ trials, i.e., newly randomly generated search displays. The first 8 blocks contained 20 Old trials.

Blocks 9–22 contained 10 Old displays included as-is, with the remaining 10 Old displays modified to yield the ‘Target violation’ and ‘Distractor violation’ conditions (5 trials per violation condition per block each). To generate a Target violation display, the target was swapped with a randomly chosen distractor, while leaving all other parameters (i.e., distractor location and rotation) untouched. To generate a Distractor violation display, the target location remained the same, but each individual distractor was rotated a random multiple of 90 ° (Figure 1B). Within each block, different Old displays were selected to be converted into a violation display, and trials were presented in random order.

After the main search task, participants were asked “Did you have the feeling that some of the search displays occurred multiple times over the course of the experiment?” and indicated their answer with a left (‘yes’) or right (‘no’) button press. Next, they were asked: “How sure are you about your answer to the previous question?” and again pressed the left (‘very sure’) or right (‘not very sure’) button. 24/36 (75 %) of participants indicated ‘not very sure’ to this confidence question, so we did not investigate this confidence rating in further detail and instead pooled responses across confidence levels.

Finally, participants performed a recognition block, consisting of the 20 Old trials and 20 newly generated New trials, randomly intermixed, and were instructed to indicate whether they thought they had seen the display during the main search task. Responses were now self-paced (unspeeded), and participants could freely move their eyes. No feedback was presented during the recognition block.

### Apparatus

Stimuli were presented using Matlab (The Mathworks, Inc., Natick, Massachussetts, United States) and custom-written scripts using the Psychophysics Toolbox (Brainard, 1997), and back-projected onto a translucent screen (48 × 27 cm) using a ProPixx projector (VPixx Technologies, Saint-Bruno, Québec, Canada) at a resolution of 1920 × 1080 pixels and a refresh rate of 120 Hz. Participants were seated at a distance of 85 cm from the screen.

Brain activity was recorded using a 275-channel axial gradiometer MEG system (VSM/CTF Systems, Coquitlam, British Columbia, Canada) in a magnetically shielded room. 5 faulty channels were disabled during the recording, leaving 270 recorded MEG channels. During the experiment, head position was monitored using three fiducial coils (nasion/left ear/right ear). Whenever participants’ head movement exceeded ∼5mm, the experiment was manually paused and the head position was shown to the participant, who would subsequently reposition (Stolk et al., 2013). Eye position and blinks were recorded using an Eyelink 1000 eye tracker (SR Research Ltd., Mississuaga, Ontario, Canada). All data were on-line low-pass filtered at 300 Hz and digitized at a sampling rate of 1200 Hz. Immediately prior to the MEG session, participants’ headshape and the location of the three fiducial coils were digitized using a Polhemus 3D tracking device (Polhemus, Colchester, Vermont, United States) in order to facilitate subsequent source analysis.

Anatomical MRI scans were acquired using a 3T MRI system (Siemens, Erlangen, Germany) and a T1-weighted MP-RAGE sequence with a GRAPPA acceleration factor of 2 (TR = 2.3 s, TE = 3.03 ms, voxel size 1 mm isotropic, 192 transversal slices, 8 ° flip angle).

### Behavioural data analysis

The main behavioural variable of interest for the main search task was reaction time (RT), with accuracy being of secondary interest. The evolution of RT and accuracy over the time course of the experiment (Figure 1C) were smoothed ± 1 blocks (i.e., the data plotted at block *n* represents the mean of blocks *n*-1, *n*, and *n*+1). Statistical assessment of the RT data was done for the *a priori* defined window of interest of blocks 9–22 (unsmoothed). RT data for trials with incorrect responses was discarded. Since RT distributions are typically heavily skewed, the RT data was log-transformed prior to any analysis to improve normality. Furthermore, since performance improved substantially over the course of the experiment (see Figure 1C), a linear trend over experiment time was fitted and removed from the (condition-pooled) log-transformed RT data per participant before performing group-wise analyses (Figures 1D, 2B, 2C, 3, 5A). Although these preprocessing steps improved the sensitivity of our analyses considerably, we note that our conclusions do not depend on them and also hold when analyzing raw RT.

### Behavioural data modelling

We performed Bayesian model comparison in order to determine whether the reaction time difference between Old and New trials exhibited a switchpoint or not. Detrended, log-transformed reaction times were modelled as drawn from a normal distribution with a given standard deviation σ, and mean μ that could be determined by one of five models (Figure 3A): No effect (no difference in mean between conditions), No switchpoint (constant difference in mean between conditions), Switchpoint (no difference in mean between conditions until a particular point in time in the experiment and a constant difference afterwards), Linear (linearly increasing difference in mean, trialwise across the experiment time), and Blockwise linear (linearly increasing difference in mean, blockwise across the experiment time). Specifically, the detrended log-RT for an individual Old or New trial *y* with index *k* was modelled according to the following prior and likelihood structure:

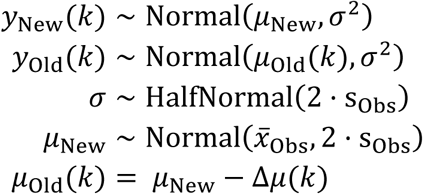

where s_Obs_ and 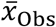 are the sample standard deviation and sample mean of the condition-pooled RT data, respectively. The No effect model was defined by Δ*μ* = 0. The No switchpoint model was defined by:

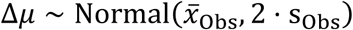

The Switchpoint model included an extra switchpoint parameter *γ* and was defined by:

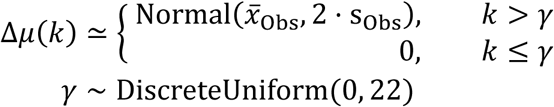

The Linear model included an extra parameter λ for the trialwise linearly increasing reaction time benefit and was defined by:

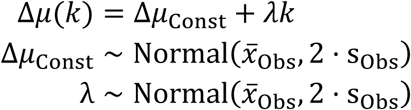

Analogously, the Blockwise linear model incorporated a blockwise linear increase:

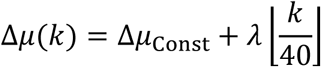

with 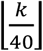 representing zero-based block index, and the other quantities as above for the Linear model.

Posterior probability distributions for the model parameters given the data were obtained per participant using Markov Chain Monte Carlo (MCMC) with a Metropolis-Hastings sampling scheme. Four chains were sampled for each model, with 40,000 samples per chain. We discarded the first 20,000 samples (burn-in), and used every 5^th^ sample of the remaining samples (i.e., thinning factor of 5). Convergence of the chains was checked by visual inspection. We computed the Watanabe-Akaike Information Criterion (WAIC) (Watanabe, 2010; Gelman et al., 2014) for each model, given the resulting MCMC posterior probability distribution. In order to compare the different models across the sample of participants, we z-scored the resulting WAIC scores across models per participant before testing. Relative WAIC scores reported in the text are expressed relative to the model with highest WAIC (i.e., the worst fitting model) per participant. Models and sampling scheme were implemented using the PyMC3 library for Python (Salvatier et al., 2016), along with Numpy (van der Walt et al., 2011) and SciPy (Jones et al., 2001).

### MEG data preprocessing and trial selection

All MEG data analyses were performed using custom-written scripts and the FieldTrip toolbox (Oostenveld et al., 2011). MEG data were segmented from −1 to +1.5 s around each search display onset. The CTF MEG system records noise in separate reference sensors, concomitantly with brain activity. As a first preprocessing step, the signals from these reference sensors were used to denoise the recorded MEG data. Next, the per-trial and per-channel mean across time was subtracted from the data. Data were screened for outliers including eye blinks or eye movements, MEG SQUID jumps, and muscle artifacts, using a semi-automatic routine (FieldTrip’s ft_rejectvisual), rejecting artifactual segments and/or excessively noisy MEG channels. This resulted in 268.9 ± 1.4 (mean ± standard deviation) MEG channels retained of the total 270 recorded, and 795.9 ± 34.5 trials per participant retained out of a total of 880. After artifact rejection, data were downsampled to 400 Hz (after applying an anti-aliasing filter) to speed up subsequent analyses. Infomax Independent Component Analysis (ICA; (Bell and Sejnowski, 1995)) was then used to clean the data of artifacts caused by ongoing cardiac activity and any residual eye movements.

### Time-frequency analysis of power

To obtain time-frequency representations of the evolution of oscillatory power over trial time (Figure 4A), we first computed an approximation of the MEG planar gradient to facilitate the interpretation of the resulting topography (Hari and Salmelin, 1997; Bastiaansen and Knösche, 2000). Power was estimated using a 500 ms sliding Hann window, centered from −0.5 to +1 s around search array onset, in steps of 50 ms, with frequency steps of 1 Hz from 1 Hz to 30 Hz. To attenuate edge artifacts, trials were zero-padded to a length of 3 s before spectral estimation. After estimating single-trial power, the two orientations of the planar gradient were combined by summing. Subsequently, we regressed out a linear trend across the experiment time from the power data to remove very slow drifts, and normalized the single-trial data by z-scoring (Grandchamp and Delorme, 2011). Power data was averaged per condition, using only those trials for which the participant responded correctly. For visualization purposes only, the time-frequency representation of the power contrast (Figure 4A) was smoothed and interpolated using a 512 × 512 grid.

### MRI preprocessing and source analysis

Anatomical MRI images were realigned to individuals’ CTF space by using the three fiducial coil locations and the recorded headshape information, using an automatic procedure followed by manual inspection and optional correction. A source model was created by non-linearly warping a regular 8 mm isotropic grid defined in Montreal Neurological Institute (MNI) space to the individual participants’ CTF space. For each grid point, the lead field was computed using a realistic single-shell volume conduction model (Nolte, 2003).

The cross-spectral density required for source analysis was estimated on the condition-pooled axial gradiometer data at the frequency of interest, as identified in the sensor-level statistical contrast (specifically: center frequency of interest 4 Hz, ± 3 Hz smoothing by using multitapers, resulting in frequency window 1–7 Hz). Beamformer spatial filters were computed from the lead field and cross-spectral density using Dynamic Imaging of Coherent Sources (DICS) (Gross et al., 2001).

We multiplied the resulting filters with the single-trial time-resolved channel-level Fourier spectra, converting the result to power values, to obtain source level estimates of time-resolved power. The source-level estimates were normalized analogously to the sensor level data, i.e. a slow linear trend across the experiment was fitted and removed and single-trial z-scoring was applied (Grandchamp and Delorme, 2011). We averaged the resulting values in the time window of interest, as identified in the sensor-level statistical contrast (0–500 ms post-stimulus), as well as per condition, and computed t-contrasts between Old and New trials across the participants. These were interpolated onto and visualized on the average MNI brain template, with scores below 65 % of the peak masked as fully transparent, everything above 80 % fully opaque, and values in between ramping up sigmoidally (Figure 4B). Note that the t-scores were computed for localization purposes only; we did not repeat the statistical test here, to avoid “double-dipping” concerns (since the effect was already deemed significant at the sensor level).

For the analysis of the evolution of the power effect over the time course of the experiment (Figure 4C), we averaged the normalized source-level data in the same 0–500 ms time window per condition, per block. For the observed hippocampus effect, the power was averaged for this single grid point of interest; while for the frontal effect, the power was averaged across the three identified peak grid points of interest (see Results for details). Analogous to the reaction time data, we increased the signal-to-noise ratio of the resulting power time courses by smoothing with ± 1 block.

### Statistical inference

For all pairwise comparisons, we report t statistics, as well as Bayes factors quantifying how much more likely the data are under the alternative hypothesis than under the null hypothesis (BF_10_). Bayes factors were estimated analytically using noninformative priors: a Cauchy prior on effect size and a Jeffreys prior on variance, using the default Cauchy scale parameter of *r* = 0.707, resulting in a quantity known in the literature as the “JZS Bayes factor” (Zellner and Siow, 1980; Jeffreys, 1998; Rouder et al., 2009).

The sensor-level mass univariate contrast (Figure 4A) was assessed using a cluster-based permutation test, which leverages the inherent correlation between the data at neighbouring time points, frequencies, and channels, to correct for multiple comparisons (Maris and Oostenveld, 2007). We used 1,000 permutations, the default maximum-sum cluster statistic, a cluster-forming threshold of 0.05 (also the default), and a minimum channel neighbour count of 2.

Error shading in figures (Figures 1C, 4C, 5A) reflects the unbiased within-participant corrected standard error of the mean (Cousineau, 2005; Morey, 2008). For visualization purposes, we furthermore highlight uncorrected p < 0.05 using thick over-or underlying bars in the same plots.

## Acknowledgments

We are grateful to Floortje Bouwkamp and Benedikt Ehinger for helpful comments on an earlier version of the manuscript. This work was supported by The Netherlands Organisation for Scientific Research (NWO Veni grant 016.Veni.198.065 awarded to ES and Vidi grant 452-13-016 awarded to FPdL) and the EC Horizon 2020 Program (ERC starting grant 678286 awarded to FPdL).

## Footnotes

1 The frequency band identified here (1–7 Hz) overlaps with the delta frequency range, but given the limited spectral resolution of the time window we are dealing with here, and the abundant prior literature on the involvement of theta activity in learning, memory, and executive function, we will label these effects as ‘theta activity’ in the rest of the paper.

## References

Addante RJ, Watrous AJ, Yonelinas AP, Ekstrom AD, Ranganath C (2011) Prestimulus theta activity predicts correct source memory retrieval. Proc Natl Acad Sci 108:10702–10707.

Awh E, Belopolsky AV, Theeuwes J (2012) Top-down versus bottom-up attentional control: a failed theoretical dichotomy. Trends Cogn Sci 16:437–443.

Backus AR, Schoffelen J-M, Szebényi S, Hanslmayr S, Doeller CF (2016) Hippocampal-Prefrontal Theta Oscillations Support Memory Integration. Curr Biol 26:450–457.

Bar M (2004) Visual objects in context. Nat Rev Neurosci 5:617–629.

Bastiaansen MC, Knösche TR (2000) Tangential derivative mapping of axial MEG applied to event-related desynchronization research. Clin Neurophysiol Off J Int Fed Clin Neurophysiol 111:1300–1305.

Bastos AM, Vezoli J, Bosman CA, Schoffelen J-M, Oostenveld R, Dowdall JR, De Weerd P, Kennedy H, Fries P (2015) Visual Areas Exert Feedforward and Feedback Influences through Distinct Frequency Channels. Neuron 85:390–401.

Bell AJ, Sejnowski TJ (1995) An information-maximization approach to blind separation and blind deconvolution. Neural Comput 7:1129–1159.

Bellmund JLS, Gärdenfors P, Moser EI, Doeller CF (2018) Navigating cognition: Spatial codes for human thinking. Science 362:eaat6766.

Biederman I (1972) Perceiving real-world scenes. Science 177:77–80.

Brainard DH (1997) The Psychophysics Toolbox. Spat Vis 10:433–436.

Brainerd CJ, Howe ML (1978) The Origins of All-or-None Learning. Child Dev 49:1028–1034.

Burgess N, Maguire EA, O’Keefe J (2002) The Human Hippocampus and Spatial and Episodic Memory. Neuron 35:625–641.

Buzsaki G, Moser EI (2013) Memory, navigation and theta rhythm in the hippocampal-entorhinal system. Nat Neurosci 16:130–138.

Chao ZC, Takaura K, Wang L, Fujii N, Dehaene S (2018) Large-Scale Cortical Networks for Hierarchical Prediction and Prediction Error in the Primate Brain. Neuron 0 Available at: https://www.cell.com/neuron/abstract/S0896-6273(18)30892-4 [Accessed November 1, 2018].

Chun MM (2000) Contextual cueing of visual attention. Trends Cogn Sci 4:170–178.

Chun MM, Jiang Y (1998) Contextual Cueing: Implicit Learning and Memory of Visual Context Guides Spatial Attention. Cognit Psychol 36:28–71.

Chun MM, Jiang Y (2003) Implicit, long-term spatial contextual memory. J Exp Psychol Learn Mem Cogn 29:224–234.

Chun MM, Phelps EA (1999) Memory deficits for implicit contextual information in amnesic subjects with hippocampal damage. Nat Neurosci 2:844–847.

Chun MM, Turk-Browne NB (2007) Interactions between attention and memory. Curr Opin Neurobiol 17:177–184.

Clark A (2013) Whatever next? Predictive brains, situated agents, and the future of cognitive science. Behav Brain Sci 36:181–204.

Cohen MA, Cavanagh P, Chun MM, Nakayama K (2012) The attentional requirements of consciousness. Trends Cogn Sci 16:411–417.

Colgin LL (2013) Mechanisms and Functions of Theta Rhythms. Annu Rev Neurosci 36:295–312.

Colgin LL (2016) Rhythms of the hippocampal network. Nat Rev Neurosci 17:239–249.

Corbetta M, Shulman GL (2002) Control of goal-directed and stimulus-driven attention in the brain. Nat Rev Neurosci 3:201–215.

Cousineau D (2005) Confidence intervals in within-subjects designs: A simpler solution to Loftus and Masson’s method. Tutor Quant Methods Psychol 1:42–45.

Dalal SS, Jerbi K, Bertrand O, Adam C, Ducorps A, Schwartz D, Garnero L, Baillet S, Martinerie J, Lachaux J-P (2013) Evidence for MEG detection of hippocampus oscillations and cortical gamma-band activity from simultaneous intracranial EEG. Epilepsy Behav 28:310–311.

de Lange FP, Heilbron M, Kok P (2018) How Do Expectations Shape Perception? Trends Cogn Sci 22:764–779.

Dehaene S, Changeux J-P, Naccache L, Sackur J, Sergent C (2006) Conscious, preconscious, and subliminal processing: a testable taxonomy. Trends Cogn Sci 10:204–211.

Eichenbaum H (2000) A cortical–hippocampal system for declarative memory. Nat Rev Neurosci 1:41.

Estes WK (1964) All-or-none processes in learning and retention. Am Psychol 19:16–25.

Fellner M-C, Volberg G, Wimber M, Goldhacker M, Greenlee MW, Hanslmayr S (2016) Spatial Mnemonic Encoding: Theta Power Decreases and Medial Temporal Lobe BOLD Increases Co-Occur during the Usage of the Method of Loci. eNeuro 3:ENEURO.0184-16.2016.

Gelman A, Hwang J, Vehtari A (2014) Understanding Predictive Information Criteria for Bayesian Models. Stat Comput 24:997–1016.

Giesbrecht B, Sy JL, Guerin SA (2013) Both memory and attention systems contribute to visual search for targets cued by implicitly learned context. Vision Res 85:80–89.

Goujon A, Didierjean A, Thorpe S (2015) Investigating implicit statistical learning mechanisms through contextual cueing. Trends Cogn Sci 19:524–533.

Grandchamp R, Delorme A (2011) Single-Trial Normalization for Event-Related Spectral Decomposition Reduces Sensitivity to Noisy Trials. Front Psychol 2 Available at: https://www.frontiersin.org/articles/10.3389/fpsyg.2011.00236/full [Accessed August 8, 2018].

Greene AJ, Gross WL, Elsinger CL, Rao SM (2007) Hippocampal differentiation without recognition: An fMRI analysis of the contextual cueing task. Learn Mem 14:548–553.

Gross J, Kujala J, Hämäläinen M, Timmermann L, Schnitzler A, Salmelin R (2001) Dynamic imaging of coherent sources: Studying neural interactions in the human brain. Proc Natl Acad Sci 98:694–699.

Hari R, Salmelin R (1997) Human cortical oscillations: a neuromagnetic view through the skull. Trends Neurosci 20:44–49.

Hasselmo ME (2005) What is the function of hippocampal theta rhythm?—Linking behavioral data to phasic properties of field potential and unit recording data. Hippocampus 15:936–949.

Hutchinson JB, Turk-Browne NB (2012) Memory-guided attention: control from multiple memory systems. Trends Cogn Sci 16:576–579.

Jeffreys SH (1998) The Theory of Probability, Third Edition. Oxford, New York: Oxford University Press.

Jensen O, Gips B, Bergmann TO, Bonnefond M (2014) Temporal coding organized by coupled alpha and gamma oscillations prioritize visual processing. Trends Neurosci 37:357–369.

Jiang Y, Wagner LC (2004) What is learned in spatial contextual cuing— configuration or individual locations? Percept Psychophys 66:454–463.

Jones E, Oliphant T, Peterson P, others (2001) SciPy: Open source scientific tools for Python. Available at: http://www.scipy.org/.

Jones JE (1962) All-or-none versus incremental learning. Psychol Rev 69:156–160.

Kahana MJ, Seelig D, Madsen JR (2001) Theta returns. Curr Opin Neurobiol 11:739–744.

Kahneman D (2013) Thinking, Fast and Slow, 1st edition. New York: Farrar, Straus and Giroux.

Kerkoerle T van, Self MW, Dagnino B, Gariel-Mathis M-A, Poort J, Togt C van der, Roelfsema PR (2014) Alpha and gamma oscillations characterize feedback and feedforward processing in monkey visual cortex. Proc Natl Acad Sci 111:14332–14341.

Koch C, Tsuchiya N (2007) Attention and consciousness: two distinct brain processes. Trends Cogn Sci 11:16–22.

Lisman JE, Jensen O (2013) The θ-γ neural code. Neuron 77:1002–1016.

Manns JR, Squire LR (2001) Perceptual learning, awareness, and the hippocampus. Hippocampus 11:776–782.

Maris E, Oostenveld R (2007) Nonparametric statistical testing of EEG- and MEG-data. J Neurosci Methods 164:177–190.

Michalareas G, Vezoli J, van Pelt S, Schoffelen J-M, Kennedy H, Fries P (2016) Alpha-Beta and Gamma Rhythms Subserve Feedback and Feedforward Influences among Human Visual Cortical Areas. Neuron 89:384–397.

Morey RD (2008) Confidence intervals from normalized data: A correction to Cousineau (2005). Tutor Quant Methods Psychol 4:61–64.

Mühlenen AV, Lleras A (2004) Spatial context and top-down strategies in visual search. Spat Vis 17:465–482.

Murthy VN (1998) Synaptic plasticity: Step-wise strengthening. Curr Biol 8:R650–R653.

Nolte G (2003) The magnetic lead field theorem in the quasi-static approximation and its use for magnetoencephalography forward calculation in realistic volume conductors. Phys Med Biol 48:3637–3652.

O’Keefe J, Nadel L (1978) The Hippocampus as a Cognitive Map. Oxford University Press.

O’Keefe J, Recce ML (1993) Phase relationship between hippocampal place units and the EEG theta rhythm. Hippocampus 3:317–330.

Oliva A, Torralba A (2007) The role of context in object recognition. Trends Cogn Sci 11:520–527.

Oostenveld R, Fries P, Maris E, Schoffelen J-M (2011) FieldTrip: open source software for advanced analysis of MEG, EEG, and invasive electrophysiological data. Comput Intell Neurosci 2011:1–9.

Osipova D, Takashima A, Oostenveld R, Fernández G, Maris E, Jensen O (2006) Theta and Gamma Oscillations Predict Encoding and Retrieval of Declarative Memory. J Neurosci 26:7523–7531.

Petersen CCH, Malenka RC, Nicoll RA, Hopfield JJ (1998) All-or-none potentiation at CA3-CA1 synapses. Proc Natl Acad Sci 95:4732–4737.

Pizzo F, Roehri N, Villalon SM, Trébuchon A, Chen S, Lagarde S, Carron R, Gavaret M, Giusiano B, McGonigal A, Bartolomei F, Badier JM, Bénar CG (2019) Deep brain activities can be detected with magnetoencephalography. Nat Commun 10:971.

Pollmann S, Manginelli AA (2009) Early implicit contextual change detection in anterior prefrontal cortex. Brain Res 1263:87–92.

Pu Y, Cheyne DO, Cornwell BR, Johnson BW (2018) Non-invasive Investigation of Human Hippocampal Rhythms Using Magnetoencephalography: A Review. Front Neurosci 12 Available at: https://www.ncbi.nlm.nih.gov/pmc/articles/PMC5932174/ [Accessed February 13, 2019].

Rouder JN, Speckman PL, Sun D, Morey RD, Iverson G (2009) Bayesian t tests for accepting and rejecting the null hypothesis. Psychon Bull Rev 16:225–237.

Salvatier J, Wiecki TV, Fonnesbeck C (2016) Probabilistic programming in Python using PyMC3. PeerJ Comput Sci 2:e55.

Spaak E, Fonken Y, Jensen O, de Lange FP (2016) The Neural Mechanisms of Prediction in Visual Search. Cereb Cortex 26:4327–4336.

Staudigl T, Hanslmayr S (2013) Theta Oscillations at Encoding Mediate the Context-Dependent Nature of Human Episodic Memory. Curr Biol 23:1101–1106.

Stolk A, Todorovic A, Schoffelen J-M, Oostenveld R (2013) Online and offline tools for head movement compensation in MEG. NeuroImage 68:39–48.

Summerfield C, de Lange FP (2014) Expectation in perceptual decision making: neural and computational mechanisms. Nat Rev Neurosci 15:745–756.

Turk-Browne NB, Scholl BJ, Johnson MK, Chun MM (2010) Implicit Perceptual Anticipation Triggered by Statistical Learning. J Neurosci 30:11177–11187.

Vadillo MA, Konstantinidis E, Shanks DR (2016) Underpowered samples, false negatives, and unconscious learning. Psychon Bull Rev 23:87–102.

van der Walt S, Colbert SC, Varoquaux G (2011) The NumPy Array: A Structure for Efficient Numerical Computation. Comput Sci Eng 13:22–30.

Watanabe S (2010) Asymptotic Equivalence of Bayes Cross Validation and Widely Applicable Information Criterion in Singular Learning Theory. J Mach Learn Res 11:3571–3594.

Zellner A, Siow A (1980) Posterior odds ratios for selected regression hypotheses. Trab Estad Investig Oper 31:585–603.

Zhao G, Liu Q, Jiao J, Zhou P, Li H, Sun H (2012) Dual-state modulation of the contextual cueing effect: Evidence from eye movement recordings. J Vis 12:11–11.

Zhao J, Al-Aidroos N, Turk-Browne NB (2013) Attention Is Spontaneously Biased Toward Regularities. Psychol Sci 24:667–677.

